# Hydrogel encapsulation of living organisms for long-term microscopy

**DOI:** 10.1101/194282

**Authors:** Kyra Burnett, Eric Edsinger, Dirk R. Albrecht

## Abstract

Imaging living organisms at high spatial resolution requires effective and innocuous immobilization. Long-term imaging, across development or behavioral states, places further demands on sample mounting with minimal perturbation of the organism. Here we present a simple and inexpensive method for rapid encapsulation of small animals of any developmental stage within a photocrosslinked polyethylene glycol (PEG) hydrogel, gently restricting movement within their confined spaces. Immobilized animals maintained a normal, uncompressed morphology in a hydrated environment and could be exposed to different aqueous chemicals. We focus in particular on the nematode *C. elegans*, an organism that is typically viewed with paralyzing reagents, nanobeads, adhesives, or microfluidic traps. The hydrogel is optically clear, non-autofluorescent, and nearly index-matched with water for use with light-sheet microscopy. We captured volumetric images of optogenetically-stimulated responses in multiple sensory neurons over 14 hours using a diSPIM light-sheet microscope, and immobilized worms were recoverable and viable after 24 hours encapsulation. We further imaged living pygmy squid hatchlings to demonstrate size scalability, characterized immobilization quality for various crosslinking parameters and identified paralytic-free conditions suitable for high-resolution single cell imaging. PEG hydrogel encapsulation enables continuous observation for hours of small living organisms, from yeast to zebrafish, and is compatible with multiple microscope mounting geometries.

## INTRODUCTION

Fluorescence microscopy has had a profound impact on biomedical research, providing spatial and temporal information about gene expression, molecular dynamics, morphology of labeled structures (Chalfie et al., 1994) and functions such as genetically encoded calcium indicators (GECIs) that indicate activity of electrically-excitable cells (Knöpfel, 2012). While many imaging experiments last only minutes per sample, long-term time-lapse microscopy can indicate changes in more gradual processes lasting hours or days. These longitudinal studies require reliable immobilization of the sample during imaging periods while simultaneously preserving the organism’s health and maintaining the function of fluorescent markers. Thus, methods of immobilization must be compatible with required environmental conditions such as hydration, temperature, and nutrition, and with imaging parameters for maximal signal with minimal phototoxicity and sample perturbation (Laissue et al., 2017).

Small model organisms are widely used for *in vivo* studies of basic physiology and systemic responses. The nematode *Caenorhabditis elegans* is a particularly useful model organism due to its <1 mm size, optical transparency, short life cycle, and ease of genetic manipulation. Standard methods for immobilization of *C. elegans* include treating the organism with paralyzing reagents such as the mitochondrial inhibitor sodium azide or the acetylcholine agonists tetramisole or levamisole (Shaham, 2006). However, chemical paralytics have disadvantages: azide causes gradual fluorophore bleaching, whereas tetramisole contracts the body, altering some physical structures. Additionally, these anesthetic reagents have toxic effects on the worm when applied during long-term studies. Another method mechanically immobilizes worms with nanobeads (Kim et al., 2013) and is advantageous for studies in which recovery and long-term animal health post-imaging are needed, such as after laser axotomy or laser cell ablation (Fang-Yen et al., 2012). In both chemical and nanobead immobilization, the animal remains in a closed environment sandwiched between a cover slip and an agar pad throughout the duration of the experiment, preventing the application of external probes (such as a microinjection needle) or chemical stimuli for neurosensory or physiological studies. Further, the closed environment limits gas exchange and leads to hypoxic conditions within minutes (Jang et al., 2016). Microfluidic traps can immobilize animals in small confined geometries for neural recordings (Chronis et al., 2007), parallel animal imaging (Hulme et al., 2007), and worm sorting applications (Aubry et al., 2015), and some have the ability to present chemicals quickly and precisely (Chronis et al., 2007; Larsch et al., 2013). For worm immobilization in physically accessible and open environments, animals have been glued for electrophysiology (Goodman et al., 2012) or mounted under paraffin oil for microinjection (Evans (ed.), 2006). In both cases, animal health can deteriorate over time and the immobilization methods are not compatible with long-term studies.

An alternative approach to immobilization encapsulates the sample in a three-dimensional hydrogel. Low-melting-point agarose is used to immobilize larger samples such as zebrafish (Renaud et al., 2011), but its relative softness allows smaller organisms to move and burrow and image quality is affected by light scattering and weak autofluorescence. A thermoreversible Pluronic hydrogel can allow periodic cycles of *C. elegans* immobilization and release, gelling at 25 °C and solubilizing upon cooling to 22 °C (Hwang et al., 2014). These hydrogels are also soft, gelation is slow, and precise temperature control is required, making them challenging to use for continuous long-term, high-resolution imaging. Alternatively, covalently-crosslinked hydrogels, such as those based on poly(ethylene-glycol) (PEG), can maintain permanent stiffness and form a gel quickly at any temperature. PEG hydrogels have been studied extensively for cell culture and other applications (Durst et al., 2011; Hou et al., 2010; Lin and Anseth, 2011) and can embed cells for long periods up to several weeks (Albrecht et al., 2006; Skaalure et al., 2015). PEG hydrogels are particularly attractive biomaterials for their tunable mechanical, diffusive, and optical properties that can be varied easily by monomer chain length and concentration (Bryant and Anseth, 2002; Choi et al., 2013; Mellott et al., 2001).

Here, we explore the use of PEG hydrogels as an embedding medium for continuous long-term imaging. In particular, we have developed a rapid, convenient method for mounting and immobilizing *C. elegans* that gently encapsulates the worm in a non-toxic, photosensitive PEG hydrogel, trapping it within a small confined volume. The covalently-crosslinked hydrogel immobilizes worms within seconds of light exposure, holds them permanently for long-term studies, yet can be easily broken to recover animals. The encapsulation process uses standard lab equipment and readily-available materials, works with any size organisms, including all larval and adult stages, and can occur at any desired temperature, unlike thermally-gelling hydrogels that require heating or cooling. The method is versatile, compatible with a wide range of hydrogel size, stiffness and diffusivity, and the degree of animal constraint can be controlled. Here, we characterize the speed and quality of immobilization for high-resolution imaging under varying conditions including light sources, substrates, polymer concentrations, and buffer conditions. The flexibility of worms allows some movement within their confined spaces, and small-scale movements can be further limited by temporary introduction of paralysis reagents, or by crosslinking under hyper-osmotic conditions without any paralytic chemicals.

Additionally, we show here that PEG hydrogel encapsulation is ideally suited for light sheet fluorescence microscopy (LSFM), an attractive imaging modality for continuous long-term 3D imaging because it reduces photobleaching by more than an order of magnitude compared with confocal systems (Y. Wu et al., 2013). LSFM has enabled monitoring of embryo development over several hours in liquid-filled open-top chambers (Ardiel et al., 2017; Kumar et al., 2014; Rieckher et al., 2015). However, diffraction-limited optical imaging occurs only when both the excitation light sheet and the emitted fluorescence pass through refractive index-matched materials, making LSFM challenging to use with standard methods for mounting larval and adult animals (McGorty et al., 2015; Rieckher et al., 2015), but convenient to use with hydrogels that are nearly index-matched with water.

Overall, PEG hydrogel encapsulation is a rapid, gentle, versatile, and inexpensive alternative mounting method useful for continuous long-term imaging, in an open format that may benefit other *C. elegans* techniques such as laser ablation and microinjection, and scalable to other model organisms such as *Drosophila* and zebrafish.

## RESULTS

### Immobilization of *C. elegans* within PEG hydrogels

Encapsulation of animals in a PEG hydrogel comprises the following steps: preparing a glass slide base and cover, placing spacers that determine hydrogel geometry onto the glass slide base, pipetting the hydrogel precursor solution, picking animals into the precursor droplet, covering the droplet with a cover slip, and crosslinking the hydrogel by brief exposure to light (**Fig. 1A**). During light exposure, the hydrogel increases viscosity until gelation up to the surface of the embedded worm (**Supplementary Video 1**). After crosslinking, the animals were fully encapsulated in the hydrogel, preventing large movements beyond their encapsulated space (**Fig. 1B**). Encapsulation occurred within seconds and could last for days, as animals could not escape. The flexibility of animals and the hydrogel enabled some movement within this confined space during contraction of body wall muscles, mostly anterior/posterior translation along the body axis and less frequently axial rotation (**Supplementary Video 2**). Gravid adults could also lay eggs into the encapsulation space. Differential interference contrast (DIC) images at 100x magnification showed minimal change in cellular morphology in animals embedded in a 10% PEG hydrogel.

**Figure 1.**
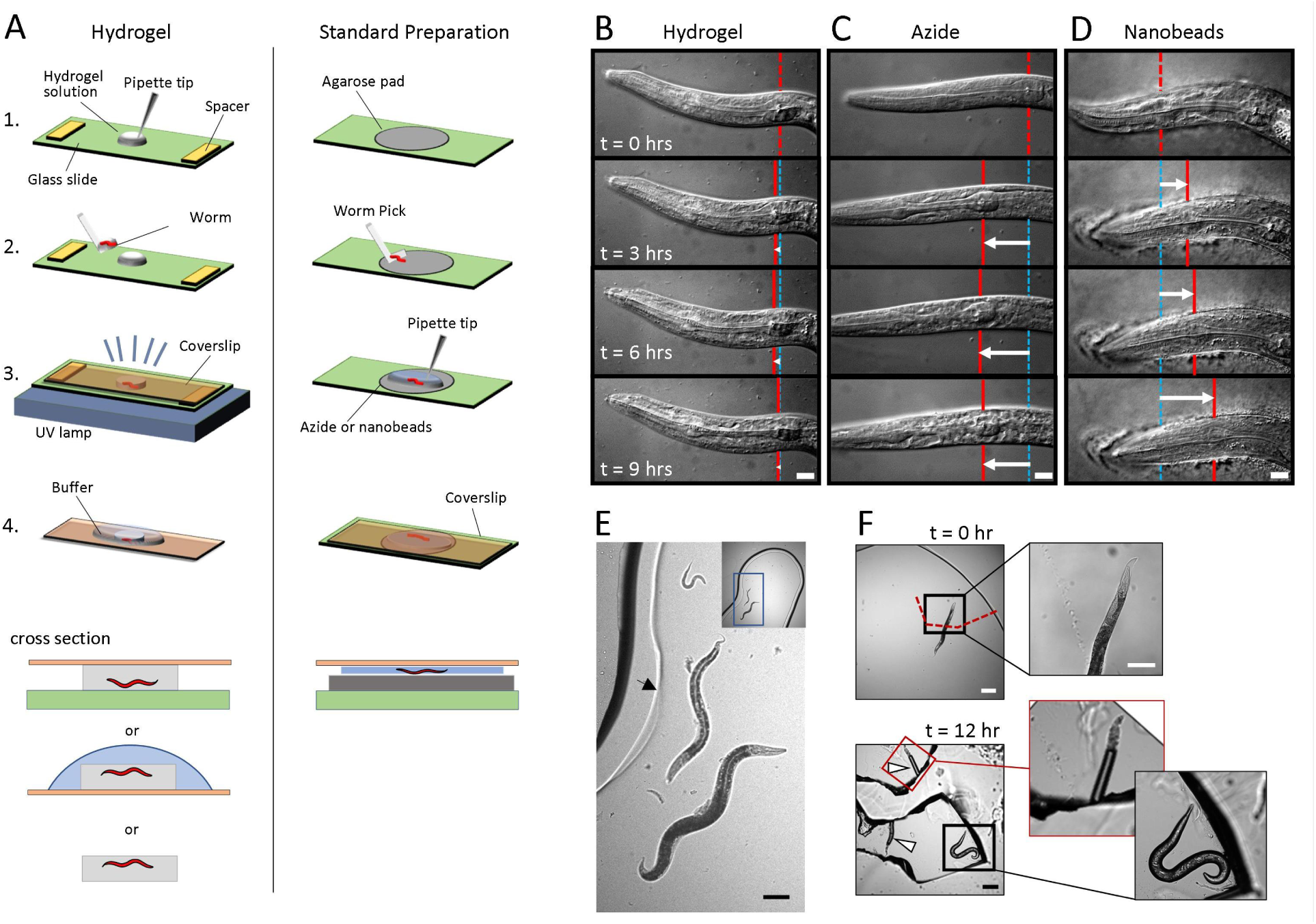
Mounting live *C. elegans* in hydrogels and on agarose pads for microscopy. (A) Schematic of worm mounting procedures by PEG hydrogel encapsulation (left) or on agar pads. (B-D) Images taken over 9 hrs in a 10% PEG hydrogel (B) or on agarose pads with 25 mM sodium azide (C) or 100 nm polystyrene beads (D). Red vertical line indicates pharynx landmark position at t = 0, blue lines indicate landmark position at each time point, and white arrow indicates displacement relative to t = 0. Scale bar, 10 μm. (E) Multiple larval and adults stages embedded in the same hydrogel. The hydrogel edge is indicated by a faint line (arrow), surrounded by about 100-200 um of uncrosslinked polymer. Scale bar, 100 μm. (F) Worms embedded in 20% PEG hydrogel were imaged immediately after crosslinking and after release 12 hr later. Arrowheads indicate the cavity of the worm in the hydrogel. Scale bars, 200 μm and 100 μm (inset).

By comparison, conventional worm mounting requires melting agarose into a thin pad, picking animals onto the pad, adding a chemical or mechanical immobilizer, and covering with a cover slip (**Fig. 1A**). Animals exposed to 25 mM sodium azide, which inhibits mitochondrial function and thereby relaxes muscle tone, can take an hour or more to fully immobilize ( **Fig. 1C**). Further, sodium azide-paralyzed animals displayed characteristics of necrotic cell death after 6 hours (Crook et al., 2013). Animals became immobilized with 100 nm polystyrene nanobeads more quickly than with azide, but they could still move gradually over hours ( **Fig. 1D**). For effective nanobead immobilization, animals were compressed causing an apparent increase the width of the worm (Kim et al., 2013).

Animals of all stages, from eggs to larval stages to adults, could be mounted in the same hydrogel (**Fig. 1E**). Encapsulated animals were recoverable by breaking the hydrogel with gentle pressure from a worm pick or fine-point forceps (**Fig. 1F**). After 24 hours of hydrogel encapsulation, young adult *C. elegans* remained mostly viable; of 241 worms, 207 (86%) crawled away upon release.

### Parameters affecting PEG hydrogel crosslinking

The mechanical, diffusive, and optical properties of PEG hydrogels can be tuned via monomer chain length and concentration (Bryant and Anseth, 2002; Choi et al., 2013; Mellott et al., 2001). To determine how gelation properties are affected by polymer concentration and other photocrosslinking parameters, we imaged animal movement during crosslinking and measured the exposure time at which motion ceased. Using several UV light sources, hydrogels of different size, monomer concentration, and photoinitiator concentration gelled in less than one minute with an irradiance dose of 15 – 220 mJ/cm^2^ (**Fig. 2A**). Higher concentrations of PEG-DA reduced the required time of UV light exposure, with a 20% hydrogel gelling in about one-half the exposure time required for a 10% hydrogel.

**Figure 2.**
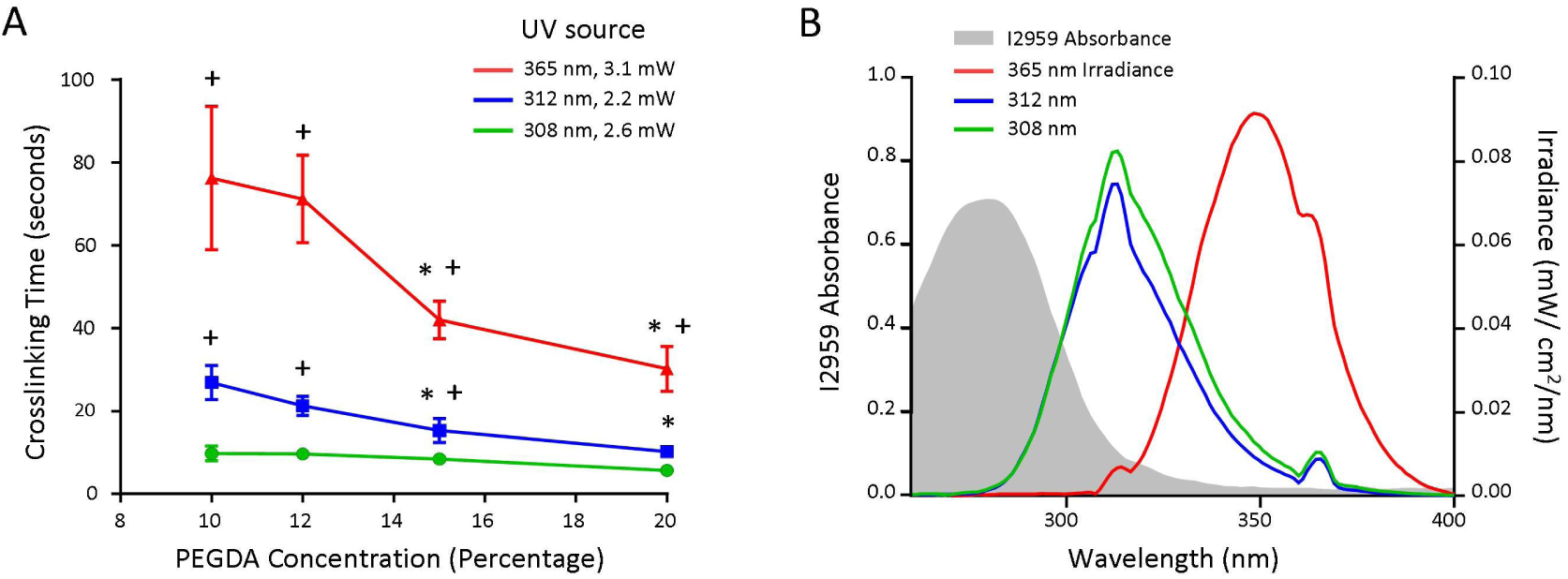
Characterization of PEG hydrogel crosslinking rates at PEG-DA concentrations of 10%, 12%, 15% and 20%. (A) Crosslinking time determined by immobilization of young adult animals in 10 - 20% PEG-DA exposed with 365 nm, 312 nm, and 308 nm UV sources. Each point represents n = 10 independent trials, with each trial having n = 2-5 worms per experiments. Values represent means (standard deviation). Statistics were performed using ordinary 2-way ANOVA with Bonferroni’s *post hoc* tests for pairwise comparisons: \**P* < 0.05 for each concentration compared to 10% PEG-DA, and +*P* < 0.05 for each source compared to 308 nm UV. (B) Absorbance spectrum of 0.001% I2959 (black, left axis), and emission spectra for each UV exposure source (right axis). Emission spectra were normalized such that area under each curve matched total output power.

The intensity and wavelength of light exposure affects crosslinking rate. The absorbance of the I2959 photoinitiator drops rapidly above 300nm (**Fig. 2B**) such that shorter wavelength UV sources crosslink the hydrogel more efficiently. For example, a 308 nm medium-wave UV-B transilluminator box gelled the polymer in 6–10s at 2.6 mW/cm^2^ (16–26 mJ/cm^2^ dose) for 20%– 10% hydrogel concentration, due to strong overlap between light emission and photoinitiator absorbance. Similarly, a 312 nm handheld medium-wave UV-B light required 12–25s exposure at 2.2 mW/cm^2^ (30–55 mJ/cm^2^ dose). A 365 nm long-wave UV-A source required a longer exposure time of 30–70 s at 3.1 mW/cm^2^ (95–220 mJ/cm^2^ dose) for 20%–10% polymer concentration due to weak photoinitiator absorbance at this wavelength. Glass slides and cover slips that absorb strongly at UV-B wavelengths (**Supp. Fig. 1**), such as those made from soda-lime glass and many plastics, require extended exposure times.

Hydrogel geometry had a minor effect on crosslinking rates. Thinner hydrogels (100 μm vs. 500 μm) and smaller hydrogels (1 μL vs. 10 μL) gelled slightly faster by less than 20% ( **Supp. Fig. 2**). Photoinitiator concentration did not affect crosslinking rate.

### Effects of buffer conditions on immobilization of *C. elegans* for microscopy

Because encapsulated animals could push against the hydrogel and move or rotate slightly, we explored modifications that would reduce micron-scale movement for long-term high-resolution microscopy (**Fig. 3**). Micron-scale movement was quantified by tracking nuclei over 3 mins (**Supp. Fig. 3**) and by a Movement Index (M.I.) sensitive to changes in position, rotation, focus, and photobleaching (calculated as the relative pixel intensity difference across frames, **Supp. Fig. 4**). The range of movement of animals embedded in 20% PEG hydrogels in water averaged 39 μm over 3 minutes (16 μm – 68 μm), with an average movement index (M.I.) of 0.33 ± 0.21. Exposure to the paralytic reagents sodium azide or tetramisole significantly reduced micron-scale motion, both when applied during gelation and when applied transiently after encapsulation (18 μm, M.I. 0.07 ± 0.01, and 19 μm, M.I. 0.06 ± 0.02 respectively). Photobleaching observed when using sodium azide contributed a majority of the M.I. compared with tetramisole, as apparent in individual worm M.I. traces (**Supp. Fig. 5**).

**Figure 3.**
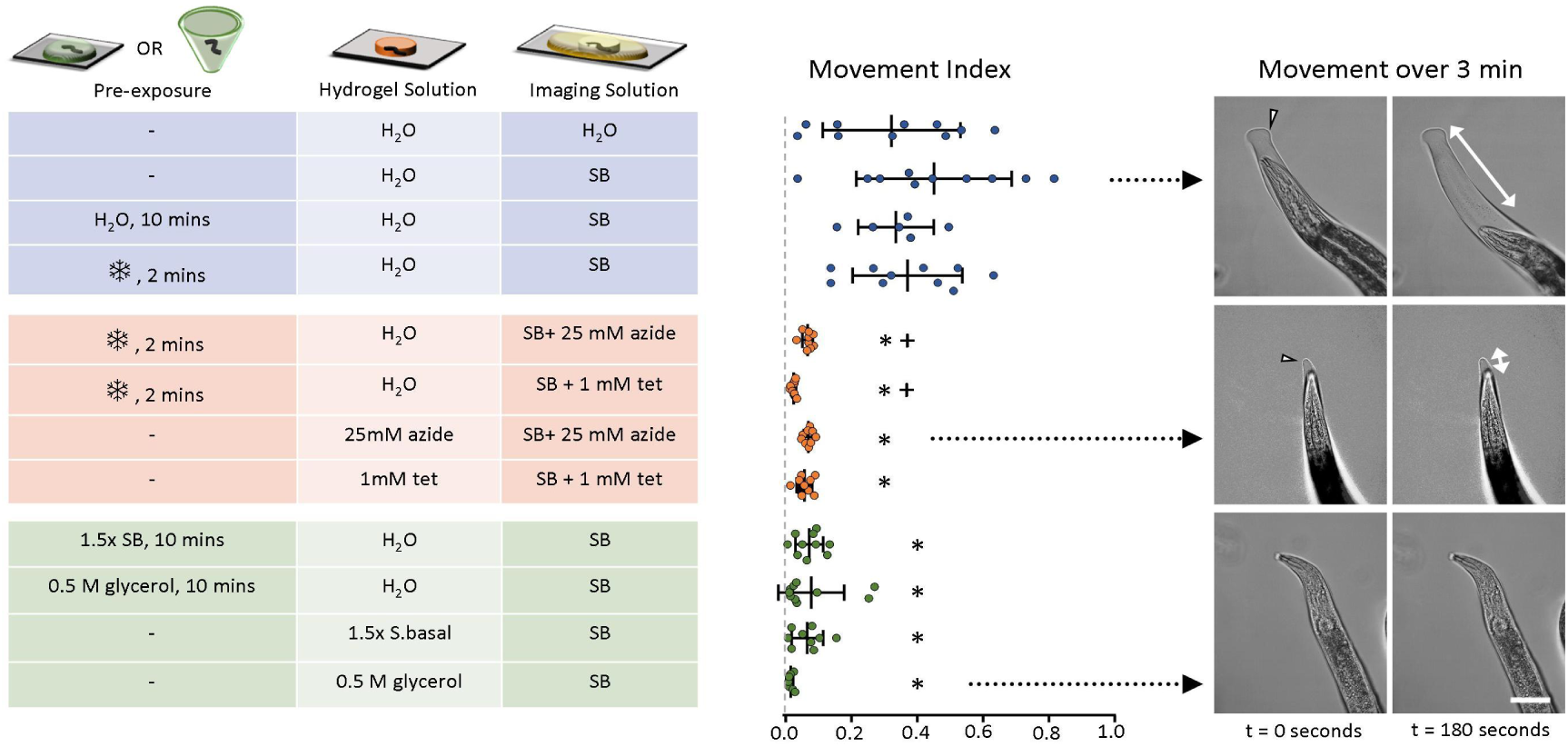
Buffer conditions before, during and after hydrogel crosslinking influence immobilization for microscopy. Pre-exposure to hypo-or hyper-osmotic solutions for 10 minutes, or cooling pretreatment (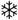 on ice or in a -20^o^C freezer for 2 minutes) occurred in a droplet or microtube. Hydrogel solutions were prepared in water (H_2_O), S-Basal buffer (SB), 25 mM sodium azide (azide) or 1 mM tetramisole (tet) in water, 500 mM glycerol in water, or 1.5x S-Basal buffer. Each dot represents the mean movement index over 3 min (Supp. Figs. 4, 5), n = 7-10 worms per condition. Vertical and error bars represent mean and standard deviation. Images represent typical movement under the conditions indicated by black arrows. Arrowheads indicate the edge of the hydrogel; animal movement can occur within this confined space (white arrows). Scale bar equals 30 μm. Statistics were performed using ordinary 2-way ANOVA with Bonferroni’s *post hoc* tests for pairwise comparisons: \**P* < 0.05 is compared with the hydrogel control with SB and +*P* < 0.05 is compared with the hydrogel cooling control with SB.

Toward paralytic-free tight immobilization, we explored the ability for cooling temperatures or osmotic changes to reduce movement. Cooling to about 4 °C on ice temporarily immobilizes animals (Chung et al., 2008). However, while cooling stopped thrashing before crosslinking, movement after embedding remained similar to uncooled animals. Cooling also allowed animals to settle within the hydrogel droplet, positioning them parallel to the glass substrate for improved imaging.

Buffer osmolarity changes body size, shrinking animals in hyper-osmotic solutions over the course of minutes through the loss of water (Lamitina et al., 2004). We reasoned that animals placed in a hyper-osmotic solution before or during gelation would shrink, thereby reducing the encapsulation space and tightening their confinement upon return to normal osmolarity. Conversely, swelling animals before crosslinking in hypo-osmotic solutions could expand the hydrogel space, thereby providing more space for movement. Crosslinking the hydrogel in water (0 mOsm), then imaging in S-Basal buffer (280 mOsm), did not significantly increase movement (39 ± 17 μm range over 3 min, M.I. 0.45 ± 0.24) compared with imaging in water (41 ± 28 μm, M.I. 0.33 ± 0.21). However, animals in a hyper-osmotic solution of 0.5 M glycerol (500 mOsm) and 1.5x S-Basal buffer (420 mOsm) for 10 minutes prior to or during crosslinking, then imaged in normal osmolarity S-Basal buffer, were well immobilized (8 ± 7 μm range, M.I. 0.02 ± 0.007 and 15 ± 9 μm range, M.I. 0.07 ± 0.05, respectively). These movement levels were comparable to immobilization by chemical paralytics.

### Imaging neural activity in encapsulated adult worms with light sheet microscopy

Hydrogels are nearly index-matched with water, therefore suitable for light sheet microscopy. We used PEG hydrogel immobilization to image young adult *C. elegans* for up to 14 hours without chemical paralytics. Worms expressing the Chrimson channel and GCaMP calcium reporter in the AWA neuron pair were imaged for three 1 hour trials over 14 hours, stimulated with 10-s pulses of red light every minute. Responses were reliably observed in the soma and neurites of both AWAL and AWAR neurons after embedding, and continued strongly in AWAR after 6 h and 13.5 h at room temperature (**Fig. 4**). Responses in AWAL declined over the first hour, and while present at the beginning of 6 h and 13.5 h trials, declined further over time. AWAR response magnitude declined about 2-fold over 14 hrs in the cell body and 5-fold in neurites.

**Figure 4.**
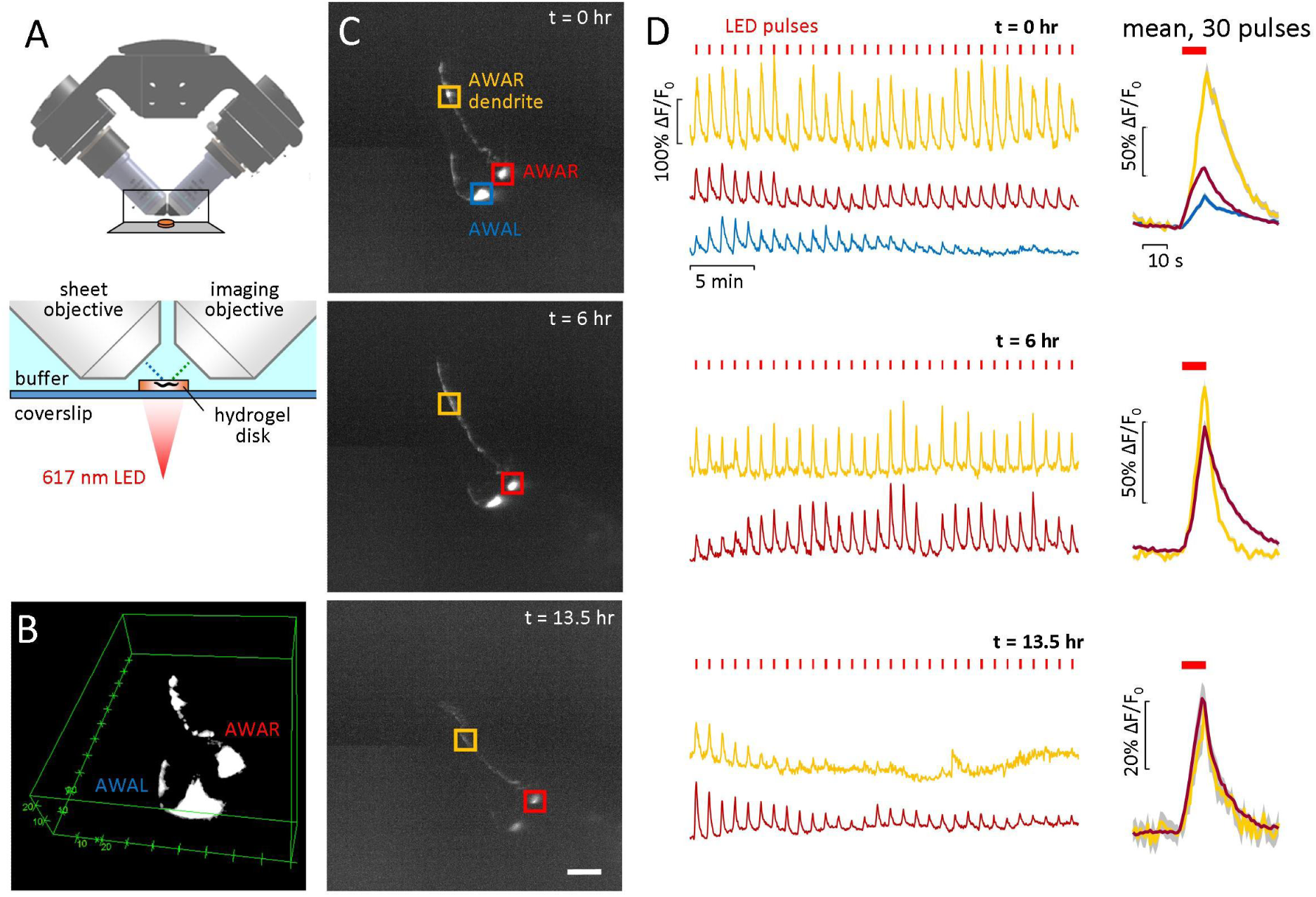
Long-term imaging of red light-activated AWA neurons expressing Chrimson and GCaMP2.2b using diSPIM. (A) Schematic of diSPIM objectives, hydrogel, and red light exposure. (B) Three dimensional volumetric view of AWA neurons. (C) Maximum intensity projection of highlighting ROIs of AWAL and AWR cell bodies and AWAR dendrite. Scale bar, 20 μm. (D) Time-lapse recording of GCaMP signal in each ROI at t=0 hr, 6hr, and 13.5 hr time points. LED was pulsed for 10 seconds, once per minute, for 1 hour; 30 mins are shown for clarity. At right, mean F/F_0_ is calculated for the 30 stimulation pulses indicated at left, with shading indicating SEM, and red bars above indicating 10-s red light exposure.

### Encapsulation and imaging of squid hatchlings

Soft mounting methods are especially important for flexible specimens, such as marine organisms. We encapsulated three-day old pygmy squid hatchlings in 1.2 mm thick, 4 mm diameter 16% PEG hydrogel disks under anaesthesia and transferred them to sea water for recovery (**Fig. 5**). After several minutes, hearts were beating and chromatophores actively opened and closed, indicating recovery of muscular behavior in internal structures. External structures in contact with the hydrogel, including arms, fins, and mantle, were immobilized. Prior to encapsulation, squid were stained with 1 uM BODIPY C3 succinimidyl ester dye to generally label cell boundaries throughout the organism (**Fig. 5C**). A volumetric stack was obtained using the diSPIM light sheet microscope through the tip of one arm of the squid ( **Fig. 5D**). Sharp cellular borders and slice alignment indicate minimal movement during image capture.

**Figure 5.**
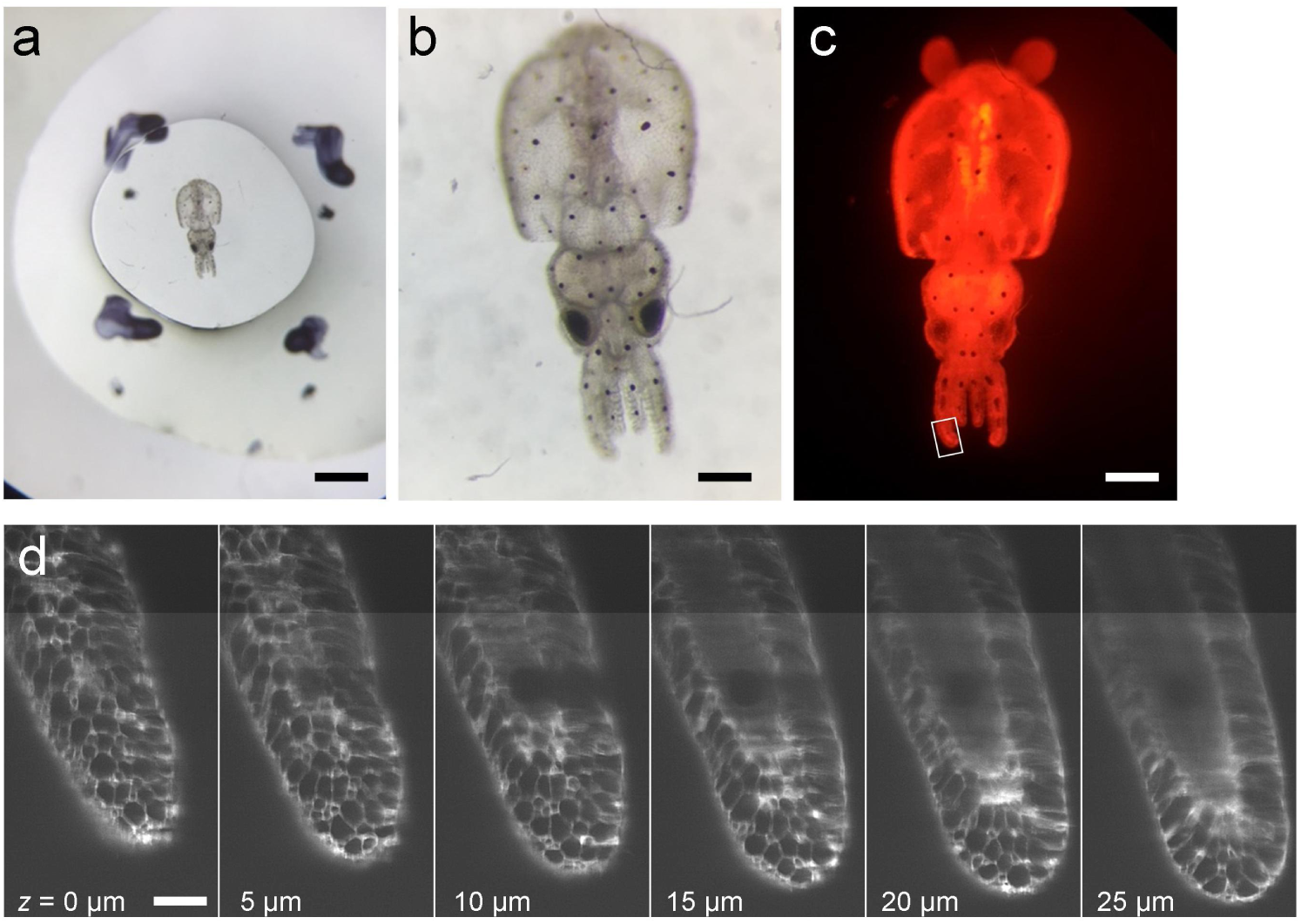
Encapsulation of a living pygmy squid hatchling in PEG hydrogel for light sheet imaging. A 3-day old squid hatchling was stained with 1 μM BODIPY C3 succinimidyl ester and embedded in a 1.2 mm thick, 4 mm diameter hydrogel disk and viewed in brightfield (a, b) and on a fluorescent dissecting microscope (c). Raw light sheet image slices of one squid arm indicated in panel c (box) are shown at 5 μm increments. Scale bars: 1 mm (a), 250 μm (b,c), 50 μm (d).

## DISCUSSION

We present here a method for gentle, effective immobilization of living organisms for long periods of time in covalently photo-crosslinked PEG hydrogels. Mounting larval and adult *C. elegans* by hydrogel encapsulation was as simple as, or potentially easier than, standard agarose preparations, and offers several advantages. The hydrogel material is inexpensive ($10/mL or 1¢/μL) and photocrosslinking equipment is already present in most biological labs. Animals were immobilized faster than chemical paralytics, more completely than with nanobeads, and without compression under glass that can alter morphology. The overall encapsulation process takes only minutes, with gelation occurring within seconds, trapping the animal within a confined space. The amount of organism movement within the hydrogel cavity could be tuned by applying different solutions that rapidly diffused into the hydrogel. For example, transient exposure to standard paralytic chemicals restricted micron-scale movements within minutes. Alternatively, we identified a paralytic-free immobilization method that temporarily reduced worm body size in hyperosmotic solutions before crosslinking, thereby tightening the hydrogel and immobilizing the animal as effectively as chemical paralytics. High-resolution image quality was maintained over 9 hours of recording in PEG hydrogels, and most animals were recoverable after 24h by physically breaking the hydrogels and allowing animals to escape from the confined space.

PEG hydrogels can be crosslinked in various sizes, making this approach suitable for embedding biological samples from <1 μm to mm or larger, a size range spanning from bacteria, yeast, and mammalian cells to small organisms including nematodes, marine organisms, *Drosphilia*, and zebrafish. We demonstrated encapsulation of living pygmy squid hatchlings, ∼1.5 mm long, whose hearts beat internally while soft motile external structures such as arms, fins, and mantle remained immobilized for light sheet or confocal microscopy.

The hydrogel mounting method is versatile and compatible with various polymer concentrations that tune mechanical stiffness and diffusive properties. Mechanical stiffness increases 10-fold from 30 to 300 kPa between 10%–20% (w/v) concentrations of PEG-diacrylate (PEG-DA, 3 kDa), whereas mesh size decreases from 4 nm to 3 nm over this range (Nguyen et al., 2012). In these hydrogels, small molecules can diffuse freely, but larger proteins cannot. Longer PEG-DA monomer chains can extend pore size over 10 nm, enabling diffusion of small proteins. Bacterial food would not permeate the hydrogel and embedded animals could not be fed by applying a becterial suspension, although they could be nourished by a chemically defined medium such as *C. elegans* Maintenance Medium (CeMM) (Szewczyk et al., 2003).

Higher PEG-DA concentration reduced crosslinking time, as reported previously (Hockaday et al., 2012), and stiffer hydrogel disks were easier to handle than weaker ones. However, animal immobilization was as good in 12% gels as in stiffer 20% gels (**Supp. Fig. 6**). This suggests that a concentration of 12% may be optimal for worm immobilization with higher diffusivity when gels needn’t be moved, while higher concentrations may be preferred if gels need to be transferred manually.

PEG-DA gelation involves activation of the photoinitiator and covalent joining of acrylate polymer chain end-groups via radical chemistry. Oxygen quenches the crosslinking reaction by scavenging the initiator radicals, slowing or preventing gelation. Thus, PEG-DA solutions exposed to air do not gel, and hydrogel disks formed between glass slides retain a ∼100 μm border of ungelled polymer where exposed to air (**Fig. 1E**). This effect may explain subtle increases in exposure requirements for taller and larger hydrogel disks, which have a larger surface exposed to air. Purging the surrounding space with an inert gas, such as nitrogen, effectively reduces this ungelled border, although this step is generally unnecessary unless a precise hydrogel geometry is needed.

PEG hydrogels can be crosslinked with numerous photoinitiators sensitive to UV or visible light. Irgacure 2959, a UV-sensitive initiator, was chosen for its rapid crosslinking time and compatibility with cell culture (Albrecht et al., 2006; Lin and Anseth, 2011). Most visible light photoinitators are either insoluble, cytotoxic, or slow to gel (Lin et al., 2013; Oliver et al., 2016), although lithium phenyl-2,4,6-trimethylbenzoylphosphinate (LAP) is a good blue-light-activated alternative (Fairbanks et al., 2009). UV exposure has the potential to cause DNA damage, especially at short wavelengths. However, the UV-B irradiance at 2–3 mW/cm ^2^ is comparable to sunlight; in fact, PEG hydrogels will crosslink with sun exposure alone. Thus, minimal DNA damage is expected during the ∼15s exposure. Wild-type animals can also detect blue and UV light via the LITE-1 channel and respond with transient avoidance behaviors (Edwards et al., 2008).

Brief exposure to photoinitiator free radicals could also cause oxidative damage. While assays for oxidative stress were not performed here, animals remained mostly viable even after 24h encapsulation, including UV and radical exposure during crosslinking, suggesting minimal physiological perturbation. In cell culture, pre-incubation with antioxidants such as ascorbate reduced effects of free radicals on cell viability (Albrecht et al., 2005; Q. Wu et al., 2013) without affecting hydrogel crosslinking; thus, any oxidative stress noted in *C. elegans* could potentially be mitigated by similar pre-exposure to antioxidants prior to crosslinking.

PEG hydrogels could be crosslinked by a variety of UV-A and UV-B lamps, many already available in biological labs for DNA gel documentation. Gelation time is dependent on the amount of light absorbed by the photoinitiator: shorter-wave UV-B sources (308, 312 nm) crosslinked faster as they better match the photoinitiator absorbance compared with UV-A sources (365 nm). A narrow-range LED flashlight (365 nm) could also crosslink the hydrogel, but required over 6 mins exposure due to minimal spectral overlap (data not shown). Shorter-wave UV can be blocked by different glass materials (**Supp. Fig. 1**). For example, UV-C lamps (254 nm), should overlap well with I2959 absorbance, but did not cause gelation as most glass and cover slips absorb nearly all UV light in this band. With UV-B sources, we found substrates composed of soda-lime glass required about twice the exposure time as borosilicate glass.

For time-lapse microscopy, embedded animals were easily identifiable across time points, as they could not escape. However, they could still move within their encapsulation volume more than might be desired for high-resolution microscopy. Paralytics could diffuse into the hydrogel, limiting movement to <8 μm over 3 min. Altering osmolarity before crosslinking could immobilize a worm to the same level without paralytics. While long-term exposure to high osmotic conditions negatively impacts animal health (Lamitina et al., 2004), the physiological consequences of transient hyper-osmotic exposure over minutes remain to be studied. Overall, the choice to immobilize by osmotic or chemical methods will depend on the biological process under investigation. For example, neural studies in which fluorescent intensity of calcium reporters is the readout are strongly impacted by the use of the paralytic 2,3-butanedione monoxime (BDM), which inhibits Type II myosin but also affects calcium channels (Petzold et al., 2011), and sodium azide, which causes steady photobleaching over tens of minutes ( **Supp. Fig. 5**). Conversely, the acetylcholine agonist tetramisole contracts muscle and alters body shape, making it suitable for neural studies but perhaps not for morphological readouts. While immobilization itself plays little role in some biological functions, such as sensory neural responses (Larsch et al., 2013), others that are influenced by body movement, such as neuromuscular activity and organismal development, may need methods that allow periodic movement between imaging sessions (Hwang et al., 2014; Keil et al., 2017). Outside of these situations, we propose that our hydrogel encapsulation method, with hyper-osmotic pre-exposure, may serve a wide variety of imaging modalities and biological applications.

Light sheet fluorescence microscopy (LSFM) can record the same sample for hours, such as the entirety of *C. elegans* embryo development (Y. Wu et al., 2013), because it requires less excitation light and reduces photobleaching and phototoxicity. However, LSFM requires that both the excitation light and the emitted fluorescence pass through refractive index-matched materials, complicating its use with standard *C. elegans* mounting methods (Rieckher et al., 2015). Most LSFM protocols embed organisms in low melting point agarose, but image quality is reduced by animal movement, light scattering, and autofluorescence. As a demonstration of long-term 3D microscopy, unparalyzed young adult animals were embedded in PEG hydrogels and imaged in a diSPIM light sheet microscope for over 14 hours. Reliable calcium transients were recorded in the soma and processes of the AWA chemosensory neurons expressing the calcium sensor GCaMP, triggered by optogenetic red-light activation of Chrimson cation channels. We observed no autofluorescence of the hydrogel itself. Neural response magnitude declined over hours, yet these results suggest overall animal viability and the ability to observe changes in functional reporters over time scales that span state changes, which may last several hours. In comparison, photobleaching typically limits spinning-disk confocal microscopy (SDCM) recordings to about 15 - 30 minutes (Kato et al., 2015; Nguyen et al., 2016; Venkatachalam et al., 2016), due to its requirement for far greater excitation intensity than LSFM. SDCM is also difficult to use with simultaneous optogenetic activation and optical readout of calcium activity, due to partial spectral overlap between light-sensitive channel and calcium sensor excitation wavelengths (at least with current sensor/channel pairs such as GCaMP/Chrimson and RCaMP/ChR2). Previously, no methods could restrain hatched *C. elegans* under the open environmental conditions required by the diSPIM, whereas embryos could be directly attached to the substrate (Christensen et al., 2015). Here, by physically preventing thrashing in larval and adult worms, the hydrogel encapsulation method represents the first opportunity to study post-embryonic processes in a light sheet system, and the only current 3D neural imaging method compatible with long-term optogenetic stimulation.

Overall, this method provides researchers with a gentle, rapid, inexpensive way to immobilize *C. elegans* and other organisms for continuous long-term experiments up to several hours, as a complement to existing sample mounting methods. It may not be suitable for experiments in which feeding is required, or for experiments requiring full movement between imaging periods. Nonetheless, by providing open fluidic access to the hydrogel and the potential for paralytic-free imaging, this method benefits long-term studies of dynamic processes in *C. elegans* and other small model organisms, such as *Drosophilia* and zebrafish, and may further improve non-imaging techniques such as laser ablation of cells (Bargmann and Avery, 1995) and microinjection of DNA (Stinchcomb et al., 1985) in which recovery of healthy organisms post-immobilization is essential.

## MATERIALS AND METHODS

### Strains and culture

Nematode strains were grown on NGM plates seeded with OP50 bacteria. *C. elegans* were imaged as young adults and synchronized by picking L4 stage worms 24 hours prior the experiment and transferring them to seeded plates. Alternatively, L1 larval stage animals were picked individually from mixed-stage plates. The following strains were used: QW1217 (*zfIs124*[P*rgef-1*::GCaMP6S]; *ofls355*[P*rab-3*::NLSRFP]), with pan-neuronal expression of nuclear-localized GCaMP and mCherry (Venkatachalam et al., 2016), and CX16573 (*kyIs587*[P*gpa-6*::GCaMP2.2b, P*unc-122*::dsRed]; *kyEx5662* [P*odr-7*::Chrimson::SL2::mCherry, P*elt-2*::mCherry]), which expresses the Chrimson ion channel and GCaMP calcium indicator in the AWA neuron pair (Larsch et al., 2015). For optogenetic stimulation, L4 stage animals were transferred to agar plates containing 50 μM all trans-retinal (Sigma-Aldrich) overnight.

*I.paradoxus* pygmy squid adults were collected from sea grass beds in Nagoya, Japan and shipped to the Marine Biological Laboratory (Woods Hole, United States), where they were maintained in aquaria for several months before dying of natural causes. Mature animals readily mated and laid egg masses, with embryos hatching after one week to produce actively swimming and hunting squid larvae. While invertebrate care is not regulated under the US Animal Welfare Act, care and use of *I.paradoxus* in this work followed its tenets, and adhered to EU regulations and guidelines on the care and use of cephalopods in research.

### Preparation of materials for hydrogel embedding

PEG hydrogel solutions were prepared by combining 10% – 20% w/v poly(ethylene glycol) diacrylate (PEG-DA, 3350 MW, ESI BIO) with 0.05% – 0.10% w/v Irgacure 2959 photoinitiator (2-hydroxy-4’-(2-hydroxyethoxy)-2-metylpropiophenone, I2959, BASF) in deionized water (diH _2_O) or 1x S-basal buffer (100 mM NaCl, 50 mM KPO_4_ buffer pH 6.0). A clean 1” x 3” glass slide (VWR Micro Slides) was rendered permanently hydrophobic by exposure to vapors of (tridecafluoro-1,1,2,2-tetrahydrooctyl) trichlorosilane (Gelest) under vacuum for 1 hour, or temporarily hydrophobic by wiping with Rain-X Glass Water Repellent. Glass slides were cleaned with ethanol and water, then dried by air gun. For covalent attachment of hydrogels to glass, #1.5 cover slips (Thermo Scientific) were silanized by coating with 3-(trimethoxysilyl)propyl methacrylate (Sigma-Aldrich) (21mM in ethanol) for 3 min, followed by ethanol wash, water rinse, and air dry. Treated glass slides can be prepared months in advance. Spacers were prepared by casting polydimethylsiloxane (PDMS, Sylgard 184, Ellsworth Adhesives) in a 1:10 (curing agent:base) ratio to thicknesses of 100, 200, and 500 μm.

### Embedding live animals in PEG hydrogel

A small volume (1 – 20 μL) of PEG hydrogel solution with photoinitiator was pipetted onto a hydrophobic glass slide flanked by two PDMS spacers whose thickness matched the desired hydrogel thickness. Animals were transferred into the hydrogel solution by worm pick and optionally cooled on ice or in a freezer to slow animal movement. A cover slip, untreated or silanized, was placed over the hydrogel droplet and supported by the spacers. The glass slide/cover slip sandwich was then placed over a UV light source and illuminated until gelation, 5 – 100 sec (depending on lamp power and hydrogel concentration). The sample was either observed immediately or the hydrogel disk was exposed by lifting the cover slip and adding a drop of aqueous solution over the disk. Hydrogel disks could be transferred to wet agar dishes to keep embedded animals hydrated.

### Mounting C. elegans for observation with differential interference contrast (DIC)

For hydrogel encapsulation, 5 μL of a 10% PEG hydrogel with 0.1% I2959 containing late L1/L2 stage animals was placed on a fluorinated glass slide with 50 μm thick tape spacers, polymerized with 312 nm UV light, and imaged on an upright Zeiss Axioskop microscope with a 100x/1.4 NA objective using DIC optics. Images were captured every hour for 12 hours without movement of the sample. For comparison, animals were mounted onto conventional agarose pads with chemical or physical restraint. In azide paralysis conditions, 1 – 5 μL drop of S-Basal buffer with 25 mM sodium azide was placed on a 1% agarose pad that also contained azide. Animals were picked into this droplet and a cover slip was placed on top before imaging. Alternatively, a 0.25 – 0.5 μL drop of polystyrene nanobeads (100nm, Polysciences, Inc.) was placed on a 10% agarose pad prior to adding animals, a cover slip, and imaging (Kim et al., 2013).

### Recovery of C. elegans from PEG hydrogels

Young adult worms were embedded in 5 μL 20% PEG hydrogels with 0.1% I2959 by crosslinking for 15 s at 312 nm on an unsilanized glass slide with 100 μm tape spacers. Hydrogels were transferred to an agar plate to keep hydrated and were stored at room temperature (22–23 °C degrees). After 24 hours, hydrogels were separated with tweezers and animals were scored for viability (movement and pharyngeal pumping). Some animals were transferred to cover slips with 1% agarose pads for imaging.

### Characterization of light sources and crosslinking conditions

Several ultra-violet (UV) light sources were used for crosslinking the hydrogel: a gel box transilluminator at 308 nm (Hoefer Scientific Instruments, model UVTM-25) and two hand-held compact UV transilluminators, at 312 nm (International Biotechnologies, Inc, model UVH, 12W) and at 365 nm (UVP, model UVGL-15, 4W). Light power was measured with a power meter (Thorlabs PM100) and 200**–**1100 nm sensor (Thorlabs S120UV) placed directly on the light source or on a glass slide or cover slip. Power values were converted to irradiance by dividing by the area of the 9.5 mm diameter sensor. Illumination spectra were obtained using a spectrometer (mut GmbH TRISTAN) with a fiber light guide (200**–**1100 nm range, 400 μm diameter). The photoinitiator absorbance was obtained with a spectrophotometer (Thermo Scientific Multiskan Spectrum).

The hydrogel crosslinking time was determined optically by monitoring movement of young adult worms during crosslinking. Gels of varying PEG-DA concentration, photoinitiator concentration, and geometry (spacer thickness and volume) were compared for each light source. Videos were captured at 1 frame/s using a Leica S6D stereoscope, converted to reflectance illumination by replacing one eyepiece with a white LED lamp, and recording via the opposite eye path with a UniBrain Fire-I 580c camera. Animal movement was analyzed by comparing frame-to-frame image differences in ImageJ. Crosslinking time was determined as the time until image difference measurements reduced to within 1 standard deviation of noise, and verified visually during observation of videos.

### Quantification of movement of hydrogel-embedded animals

Young adult QW1217 worms with pan-neuronal expression of nuclear-localized mCherry were embedded in 10% and 20% PEG hydrogels (3 μL with 100 μm spacer, 0.1% I2959, 20 s exposure using a 312 nm UV source). Worms received either no pretreatment, exposure to hypoosmotic (diH_2_O, 0 mOsm) or hyperosmotic buffers (0.5 M glycerol in diH_2_O, 500 mOsm or 1.5x S-Basal, 420 mOsm) for 10 min, or cooling in a –20^o^C freezer or in contact with ice for 1 – 3 min prior to crosslinking. PEG hydrogel solutions were prepared in diH_2_O or S-Basal buffer, and some contained the paralytic reagents 25 mM sodium azide (Massie et al., 2003) or 1 mM tetramisole hydrochloride (Sigma) (Larsch et al., 2013). Other hydrogel solutions were prepared in 500 mM glycerol or 1.5x S-Basal hyperosmotic buffers. After crosslinking, all hydrogels were submerged in an aqueous solution of either S-Basal, diH_2_0, sodium azide (25 mM) or tetramisole (1mM) for imaging. Videos were captured with a Hamamatsu Orca-Flash 4.0 camera at 1 frame/sec for 3 min on a Zeiss AxioObserver inverted epifluorescence microscope with a 20x/0.5 NA objective.

Animal movement was analyzed by comparing frame-to-frame image differences (**Supp. Fig. 4**). First, the absolute value differences between each pair of consecutive frame averaged pixel intensities were calculated. Next, the difference image stack was smoothed (average of its 3 × 3 neighborhood). To reduce the contribution of pixel noise, a value of 10 (corresponding to average pixel noise) was subtracted uniformly from each frame and negative values were set to 0. Regions of Interest (ROIs) were selected over the head (nerve ring) and ventral cord and a background region (BG). The movement index was calculated as M.I. = ΔI_ROI_/ I_ROI_ – I_BG_, where ΔI_ROI_ is the mean of difference images across the ROI, and I_ROI_ and I_BG_ are the mean intensity of the ROI and background regions for the first frame, respectively. Here, identical sequential frames would have a M.I. of zero, whereas images that have changes in position, rotation, focus, or intensity have increasing M.I. values.

Movement in the image plane was quantified by tracking cell nucleus position over 3 minutes. Individual neurons were tracked using NeuroTracker (Larsch et al., 2013) and centroid positions were used to determine the range of axial movement by the animal.

### Long-term Imaging of Optogenetically-Induced Calcium Transients Using 3D Light Sheet Microscopy

Young adult animals co-expressing Chrimson and GCaMP2.2b in the AWA chemosensory neurons were embedded in a PEG hydrogel bonded to a 24 x 50 mm methacrylate-silane-treated cover slip. Animals were picked into a 2.4 μL drop of 13.3% PEG-DA solution with 0.067% I2959 in 500 mM glycerol in diH_2_O. After 5 min, the sample was cooled on ice for 30 s and exposed to UV light for 30 s with a 308 nm handheld lamp. Cover slips were mounted into a light sheet chamber (ASI, I-3078-2450) and filled with ∼5mL diH_2_O. The dual-inverted selective plane illumination microscope (diSPIM) recorded the calcium response of AWA with a 488 nm excitation laser (Vortran Stradus VersaLase) at 1 mW power setting and a 525/50 nm emission filter. Single-view volumetric stacks (40 slices with 1 μm spacing, 166 x 166 x 40 μm ^3^), were obtained at 1 volume/s for three 60 min recording sessions beginning at t = 0 h, 6 h, and 13.5 h, for a total of 432,000 image frames. A red LED light (617 nm, Mightex, with 620/30 nm filter) was mounted either above the stage, illuminating the animals at a 45 degree angle, or from below with a 600 nm shortpass dichroic through a 4x objective, and controlled via MATLAB and an Arduino controller. Red light pulses, 10 s in duration, were repeated each minute during recordings. For each time point, the volume stack was compressed into a single maximum projection plane, and intensity values were integrated across ROIs surrounding each neuron or neuronal process. After subtracting background intensity from a nearby region, fluorescence intensity (*F*) integrated across the neuron or neurite ROI was normalized to the initial intensity averaged over 1 s (*F*_*0*_). No interference was observed in background or neural ROIs during red light exposure.

### Imaging of Living Pygmy Squid Using 3D Light Sheet Microscopy

Pygmy squid (*Idiosepius paradoxus*) at three days post-hatching were treated with 1 μM BODIPY 564/570 C_3_ succinimidyl ester vital dye (ThermoFisher D2222) in filtered seawater for 1 hr to label cell boundaries and generally visualize morphology, modifying methods for the marine worm *Platynereis dumerelli* (Steinmetz et al., 2007). Squid were washed five times in filtered sea water, then paralyzed by exposure to 3.75% MgCl_2_ in sea water for 15 minutes and transferred to a hydrophobic glass slide. Excess sea water was aspirated by pipette such that approximately 4 μL liquid remained. To this, 16 μL of 20% PEG-DA in sea water with photoinitiator was added and gently mixed. A methacrylate silane-treated 24 x 50 mm ^2^ cover slip was placed 1.2 mm over the droplet using glass slides as spacers. The hydrogel was crosslinked by exposure to 365 nm UV light for 1 min. The coverslip was mounted into the light sheet chamber and filled with sea water. Dual-view volumetric stacks (30 slices with 1 μm spacing, 332 x 332 x 30 μm^3^) were obtained using a 561 nm laser (4 mW power setting).

### Statistical Analysis

Statistical comparisons were made by two-way ANOVA with significance level set at a = 0.05, followed by Bonferroni’s multiple comparison tests. Data are presented as mean ± standard deviation, unless otherwise noted.

## ACKNOWLEDGMENTS

KB and DRA designed the experiments, analyzed the data, and wrote the paper. EE contributed reagents and wrote the paper. We thank Ross Lagoy, Daniel Lawler, Naomi Otoo, Olivia Leavitt, Laura Aurilio, Connor Haley, Chris Chute, and SoRi Jang for testing hydrogel encapsulation and providing feedback. We thank Thom Geer (Nobska Imaging), Jon Daniels (ASI), the laboratories of Hari Shroff at NIH (Evan Ardiel, Ryan Christensen, Harshad Vishwasrao, Abhishek Kumar, Yicong Wu) and Daniel Colon-Ramos at Yale (Lin Shao, Mark Moyle) for assistance with light sheet imaging, and Takeshi Kasugai (Nagoya Public Aquarium) for collecting and shipping squid. DRA is supported by NSF, NIH, and holds a Career Award at the Scientific Interface (CASI) from the Burroughs Wellcome Fund. KB is supported by an NSF IGERT award.

## SUPPLEMENTARY INFORMATION

**Supplementary Figure 1.**
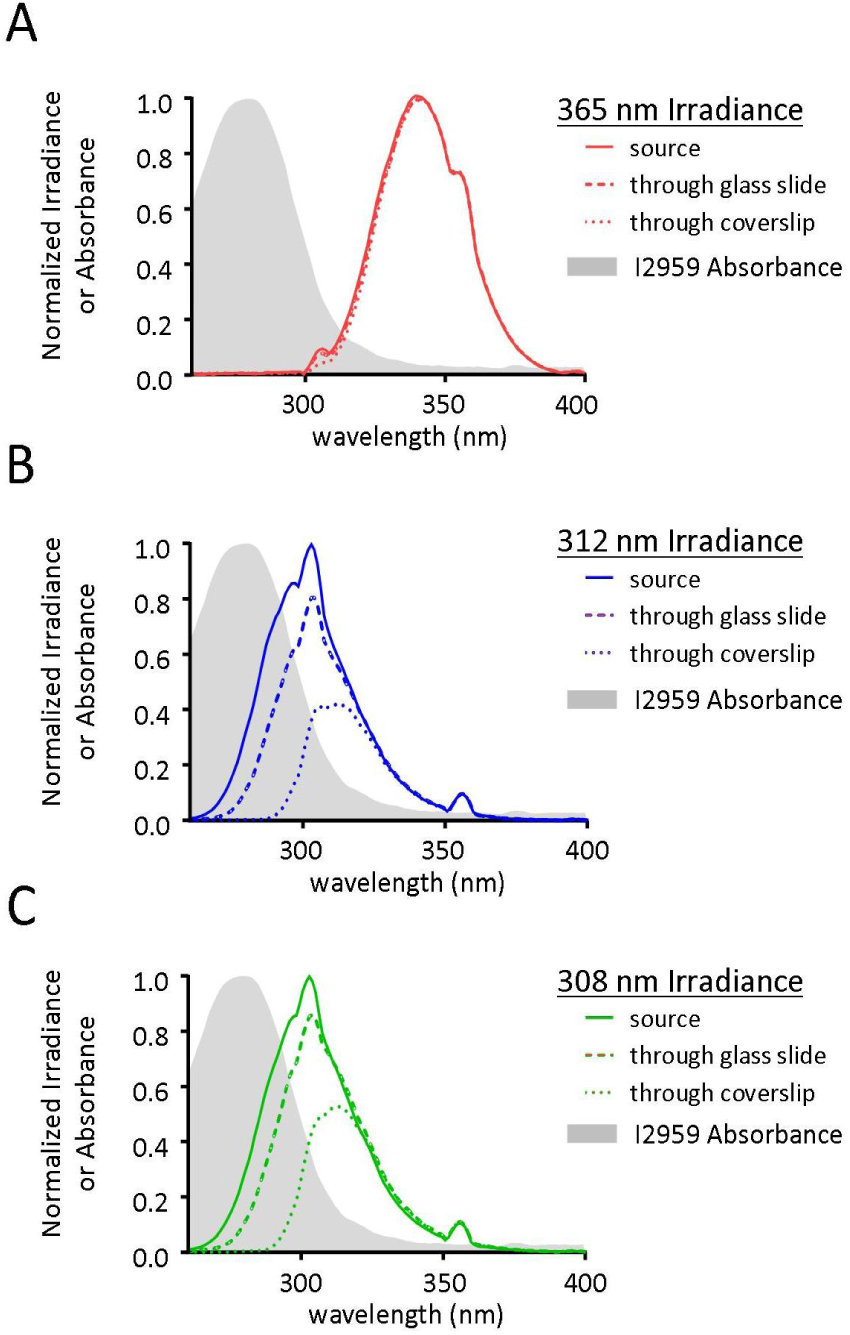
Irradiance of different UV light sources and substrates. Normalized absorbance spectrum of Irgacure 2959 photoinitiator and irradiance of each UV exposure source alone, through a glass slide, and through a glass coverslip. Irradiance curves for each substrate were normalized to the irradiance of the each source at 365 nm.

**Supplementary Figure 2.**
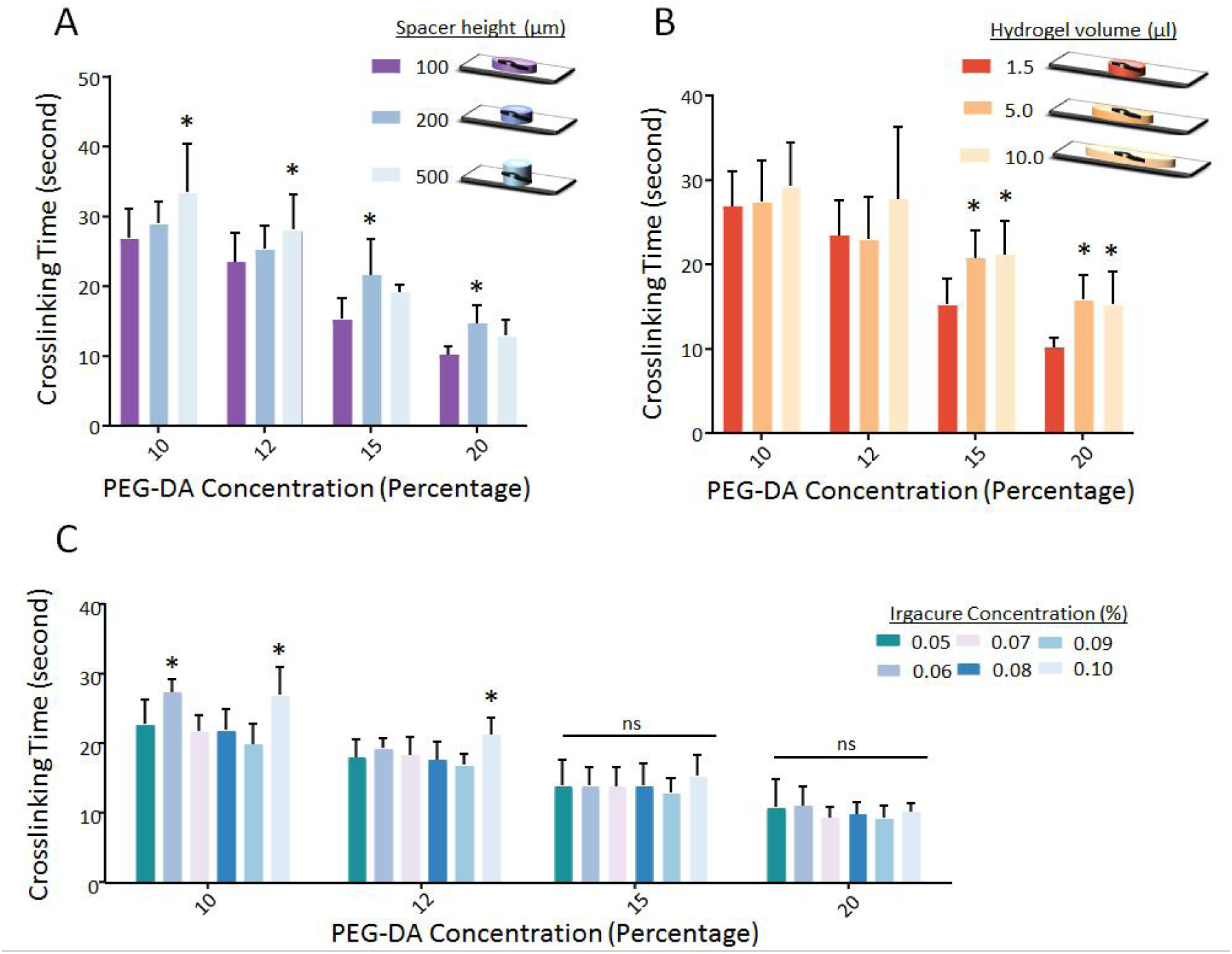
Characterization of PEG hydrogel crosslinking for different PEG-DA concentrations. Each bar represents the average amount of time it takes the hydrogel to crosslink with various volumes, heights, and photoinitiator concentrations, exposed with 312 nm UV source. (A) Spacer height varied from 100 to 500 μm, with 1.5 μL volume and 0.10% I2959. (B) Hydrogel volume varied from 1.5 to 10 μL, with 100 μm spacer and 0.10% I2959. (C) Irgacure 2959 photoinitiator concentration varied from 0.05% to 0.1%, with 1.5 μL volume and 100 μm spacer. Each bar represents n = 10 trials, with each trial averaging times from 2-5 worms. Bars represent mean and standard deviation. Statistics were performed using ordinary 2-way ANOVA with Bonferroni’s *post hoc* tests for pairwise comparisons, \**P* < 0.05 compared to 100 μm (A), 1.5 μL (B) and 0.05% (C) conditions.

**Supplementary Figure 3.**
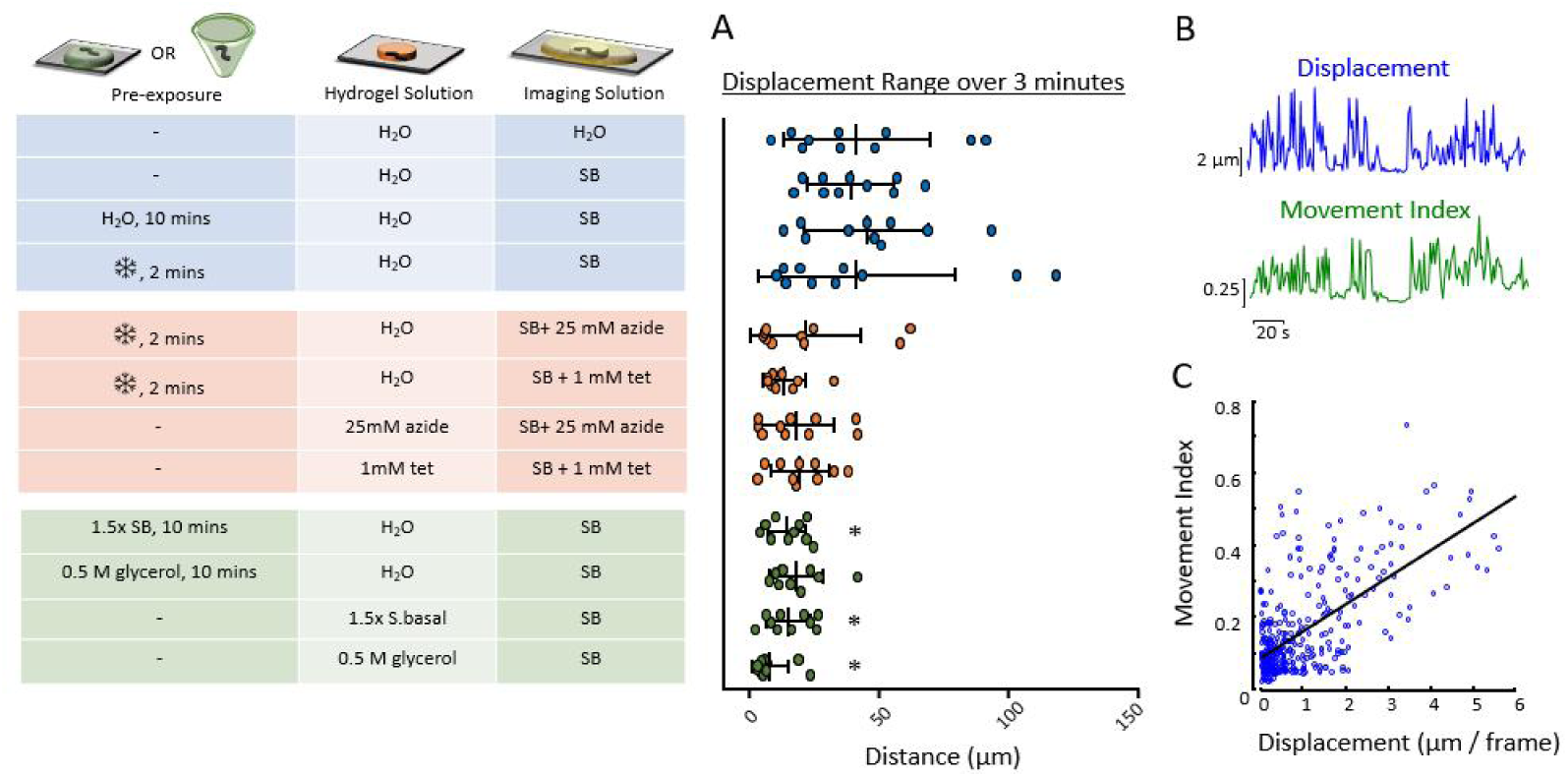
Displacement range over 3 minutes for animals embedded in different conditions before, during, and after hydrogel crosslinking. Pre-exposure to hypo-or hyper-osomotic solutions for 10 minutes, or cooling pretreatment (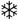 on ice or in a -20^o^C freezer for 2 minutes) occurred in a droplet or microtube. Hydrogel solutions were prepared in water (H_2_O), S-Basal buffer (SB), 25 mM sodium azide (azide) or 1 mM tetramisole (tet) in water, 500 mM glycerol in water, or 1.5x S-Basal buffer. (**A**) Mean displacement range over 3 min, from n= 7-10 worms. Vertical line and error bars represent mean (standard deviation). (**B**) Comparisons between displacement and movement index are shown for 1 worm. (**C**) Correlation of movement index versus displacement, R^2^ = 0.45. Statistics were performed using ordinary 2-way ANOVA with Bonferroni’s *post hoc* tests for pairwise comparisons: \**P* < 0.05 is compared with the hydrogel control with S-Basal solution.

**Supplementary Figure 4.**
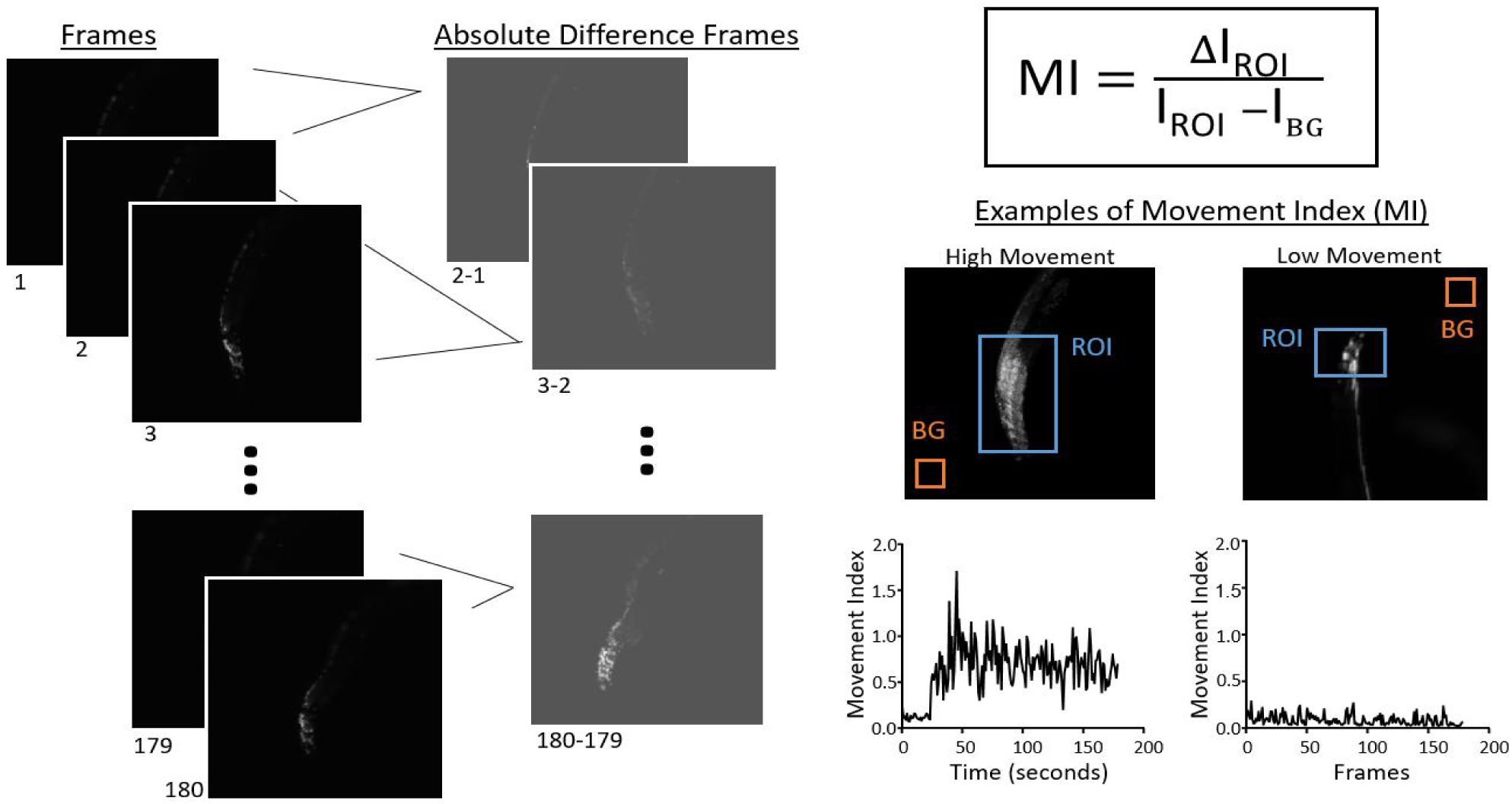
Movement Index Calculations. Movement index is described using the equation M.I. = ΔI_ROI_/ I_ROI_ – I_BG_, where ΔI_ROI_ represents the difference in pixel intensity between consecutive frames, averaged across a region of interest (ROI), I_ROI_ represents the average intensity of the ROI, and I_BG_ represents the average intensity of a Background region. High-and low-movement examples show the time course of instantaneous movement over 180 frames at 1 fps.

**Supplementary Figure 5.**
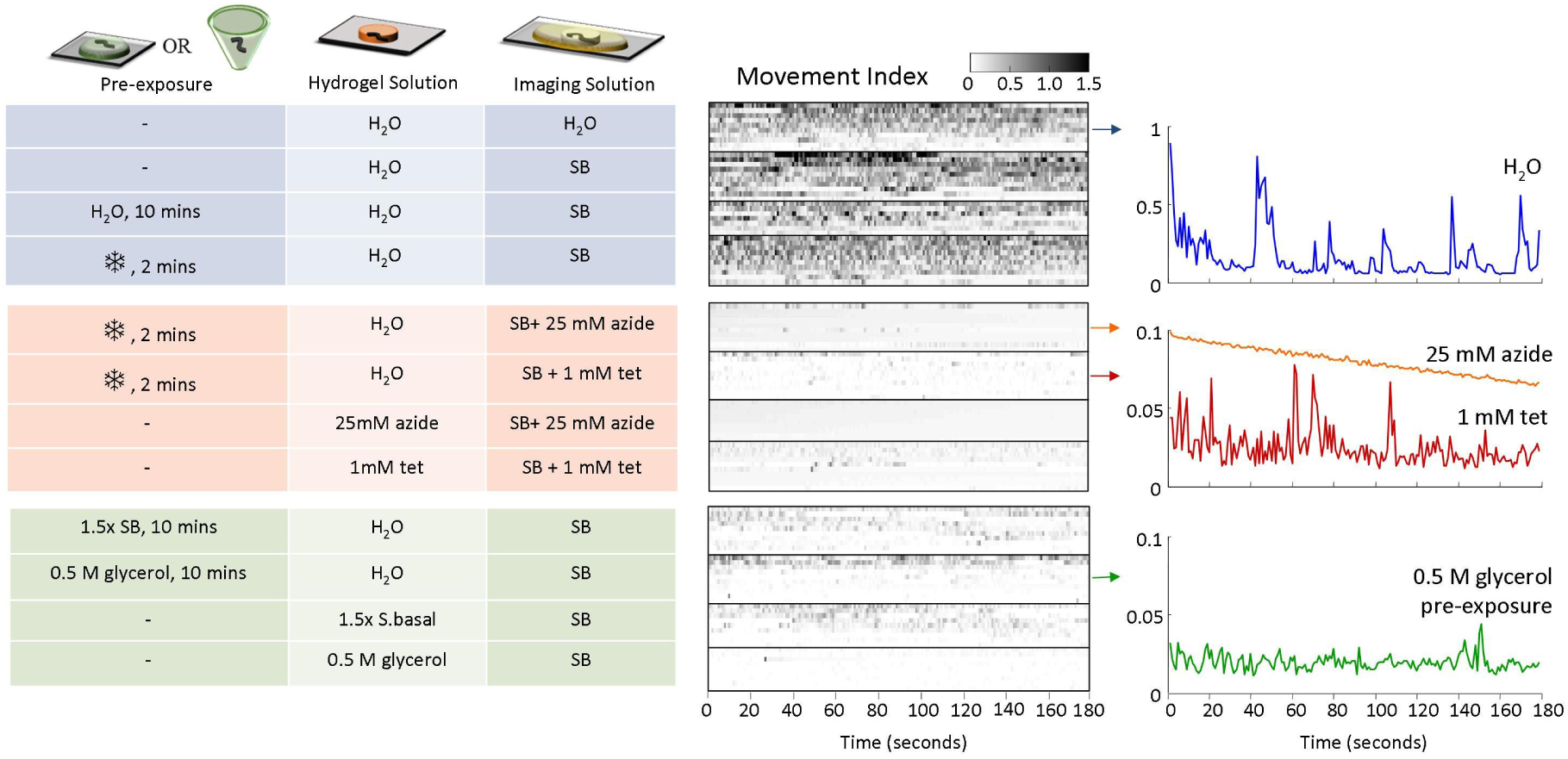
Buffer conditions before, during and after hydrogel crosslinking influence worm’s variability in movement index during microscopy. Pre-exposure to hypo-or hyper-osomotic solutions for 10 minutes, or cooling pretreatment (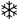 on ice or in a -20^o^C freezer for 2 minutes) occurred in a droplet or microtube. Hydrogel solutions were prepared in water (H_2_O), S-Basal buffer (SB), 25 mM sodium azide (azide) or 1 mM tetramisole (tet) in water, 500 mM glycerol in water, or 1.5x S-Basal buffer. Heat maps of movement index vs. time for the 3-minute duration of fluorescent imaging. In the middle panel, each line represents the movement index of an individual animal. On the right panel, the median movement index of each treatment group is represented.

**Supplementary Figure 6.**
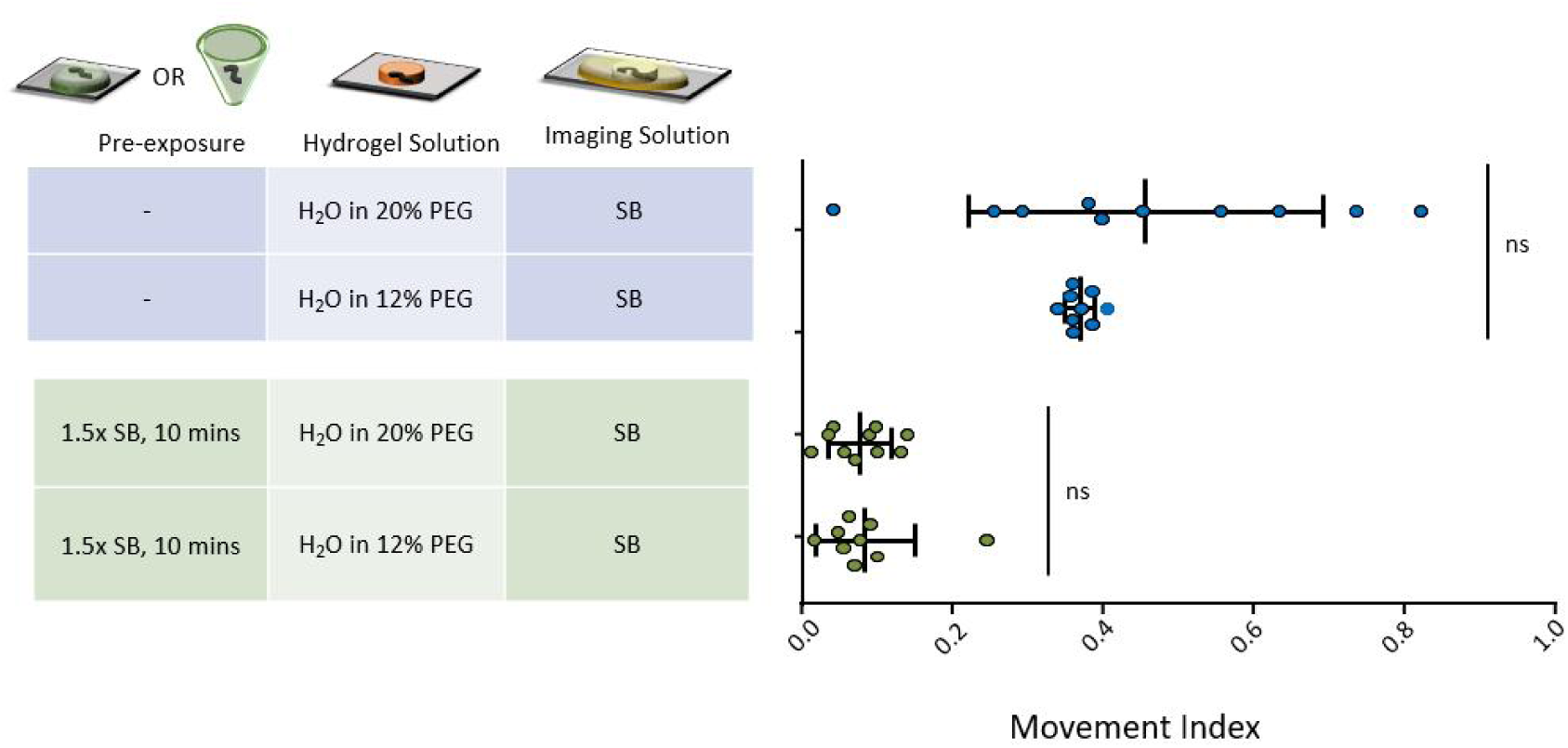
Movement of animals embedded in 20% and 12% PEG-DA hydrogels was similar under control and hyper-osmotic pre-exposure conditions (10 minutes pre-exposure to 1.5x S. Basal buffer in a droplet or microtube). Each dot plot represents the mean movement index over 3 min, from n= 9-10 worms. Vertical and error bars represent mean (standard deviation). Statistics were performed using ordinary 2-way ANOVA with Bonferroni’s *post hoc* tests for pairwise comparisons. ns, not significant.

**Supplementary Video 1.** Crosslinking of *C. elegans* in hydrogel disks. Video shows young adult animals swimming in polymer solutions of 12%, 15%, and 20% PEG-DA. At 0:06, the 312 nm UV lamp is turned on, and animals become encapsulated several seconds later. Video is accelerated 5x. Note that cooling the solution eliminates movement before gelation, allowing positioning of animals within the hydrogel disk.

**Supplementary Video 2.** Movement of young adult *C. elegans* embedded in 20% PEG-DA hydrogels. Animals were imaged in buffer (top), in 1mM sodium azide (middle), and buffer following pre-exposure to 500 mM glycerol as in Fig. 3. Video is accelerated 3x.

